# CAR Treg synergy with anti-CD154 mediates infectious tolerance to dictate heart transplant outcomes

**DOI:** 10.1101/2024.09.20.614149

**Authors:** Samarth S Durgam, Isaac Rosado-Sánchez, Dengping Yin, Madeleine Speck, Majid Mojibian, Ismail Sayin, Grace E Hynes, Maria Luisa Alegre, Megan K Levings, Anita S Chong

## Abstract

Successful allograft specific tolerance induction would eliminate the need for daily immunosuppression and improve post-transplant quality of life. Adoptive cell therapy with regulatory T cells expressing donor-specific Chimeric Antigen Receptors (CAR-Tregs) is a promising strategy, but as monotherapy, cannot prolong the survival with allografts with multiple MHC mismatches. Using an HLA-A2-transgenic haplo-mismatched heart transplantation model in immunocompetent C57Bl/6 recipients, we show that HLA-A2-specific (A2) CAR Tregs was able to synergize with low dose of anti-CD154 to enhance graft survival. Using haplo-mismatched grafts expressing the 2W-OVA transgene and tetramer-based tracking of 2W- and OVA-specific T cells, we showed that in mice with accepted grafts, A2.CAR Tregs inhibited endogenous non-A2 donor-specific T cell, B cell and antibody responses, and promoted a significant increase in endogenous FoxP3^+^Tregs with indirect donor-specificity. By contrast, in mice where A2.CAR Tregs failed to prolong graft survival, FoxP3^neg^ A2.CAR T cells preferentially accumulated in rejecting allografts and endogenous donor-specific responses were not controlled. This study therefore provides the first evidence for synergy between A2.CAR Tregs and CD154 blockade to promote infectious tolerance in immunocompetent recipients of haplo-mismatched heart grafts and defines features of A2.CAR Tregs when they fail to reshape host immunity towards allograft tolerance.

## Introduction

Inducing allograft specific tolerance would reduce the need for daily pharmacological immunosuppression and its associated side effects, as well as improve long-term allograft survival. One way to induce tolerance is combined transplantation of hematopoietic stem cells with solid organ allografts; while this has resulted in some clinical success, conditioning regimen intensity and risk of graft versus host disease limit its broad application(1–3). Another strategy is to harness the natural immunosuppressive properties of forkhead box protein P3 (FoxP3)-expressing regulatory T cells (Tregs) through adoptive cell therapy (2, 3). This approach can support the survival of multiple antigen-mismatched allografts, via the ability of Tregs to modulate antigen presenting cells and promote the development of Tregs with multiple donor specificities through a process known as infectious tolerance (4, 5). Building upon evidence in mice that infusion of polyclonal, expanded Tregs can induce transplantation tolerance (6–8), several Phase 1 trials have tested the effects of infused polyclonal Tregs in solid organ or stem cell transplantation. These studies have uniformly shown that the approach is feasible, safe, and may allow a reduction of pharmacological immunosuppression (8–11).

One limitation of cell therapy with polyclonal Tregs is that only a small proportion are alloantigen specific. Work in pre-clinical models of transplantation has shown that antigen-specific Tregs are significantly more effective than are polyclonal cells(12)^-22^. Alloantigen-specific Tregs can be generated by repetitive stimulation with donor-derived antigen-presenting cells (APCs)(13–16) or donor-antigen specificity can be introduced into Tregs by expression of Chimeric Antigen Receptors (CARs)(9, 17). Several groups have expressed HLA-A2-specific CARs (A2.CARs) in human Tregs and showed that they have superior therapeutic effects over polyclonal Tregs in humanized mouse models of graft versus host disease or skin transplantation(18, 19). These observations have led to a rapid clinical translation of this approach (NCT05234190, NCT04817774).

In order to dissect mechanistic effects and iteratively improve efficacy, A2.CAR mouse Tregs have more recently been studied in mouse transplant models and reported to delay rejection of A2-mismatched skin grafts and inhibit A2-specific antibody formation (20, 21). Similarly, in a model of A2-mismatched heart transplantation (HTx), A2.CAR Tregs significantly promoted graft survival over polyclonal Tregs, although graft palpation scores started to decline by day 20 post-HTx. However, in a haplo-mismatched A2.C57Bl/6 (B6) X BALB/c (A2.F1) model of HTx, A2.CAR Treg therapy alone was ineffective and significant extension of allograft survival was observed only when combined with 9 days of rapamycin treatment (22). Notably, this combination did not induce long-term graft acceptance, highlighting the need for further work to identify combination therapies capable of synergizing with CAR Tregs to achieve stable transplantation tolerance.

We previously reported that co-stimulation blockade with CTLA4-Ig and rapamycin prevented the expansion of both endogenous donor-specific FoxP3^neg^ CD4^+^ T conventional (Tconvs) and Tregs. On the other hand, anti-CD154 permitted Treg expansion while inhibiting Tconv expansion(23), consistent with Tregs expressing lower levels of CD154 compared to Tconvs (24). Together with the resurgence of interest in clinical use of modified versions of anti-CD154 without pro-thrombotic properties (25) (NCT04046549, NCT05983770), in this study we tested whether anti-CD154 could enhance the pro-tolerogenic properties of CAR Tregs.

We used a clinically relevant haplo-mismatched (A2.2W-OVA.F1) HTx model in immunocompetent B6 recipients and show that A2.CAR Tregs synergized with anti-CD154 to enhance graft survival and induce infectious tolerance (26). Specifically, the expansion of endogenous donor-reactive CD4^+^FoxP3^neg^ Tconvs and CD8^+^ T cells and endogenous donor-specific antibody (DSA) responses to non-A2 alloantigens were inhibited, and the expansion of endogenous donor-reactive CD4^+^FoxP3^pos^ Tregs was promoted. In contrast, when A2.CAR Tregs failed to promote graft acceptance, infectious tolerance was not observed.

## Results

### A2.CAR Tregs synergize with low dose anti-CD154 to prolong survival of haplo-mismatched heart grafts

Previous reports indicated that A2.CAR Tregs failed to promote the survival of haplo-mismatched HTx, and that adjunct immunosuppression is necessary (22). Our previous observations that anti-CD154 was superior to CTLA4-Ig or rapamycin in permitting expansion of endogenous donor-reactive Tregs (23) prompted us to test if a suboptimal dose (250 µg/mouse) of anti-CD154 could synergize with A2.CAR Tregs in B6 recipients of haplo-mismatched HTx (Figure 1A). While untreated recipients acutely rejected their heart allografts with a median graft survival of 12 days, an anti-CD154 dose of 250µg/mouse was chosen as it delayed rejection to a median survival of 28 days, whereas a higher dose of 350µg anti-CD14 uniformly induced graft acceptance (Figure 1B).

**Figure 1:**
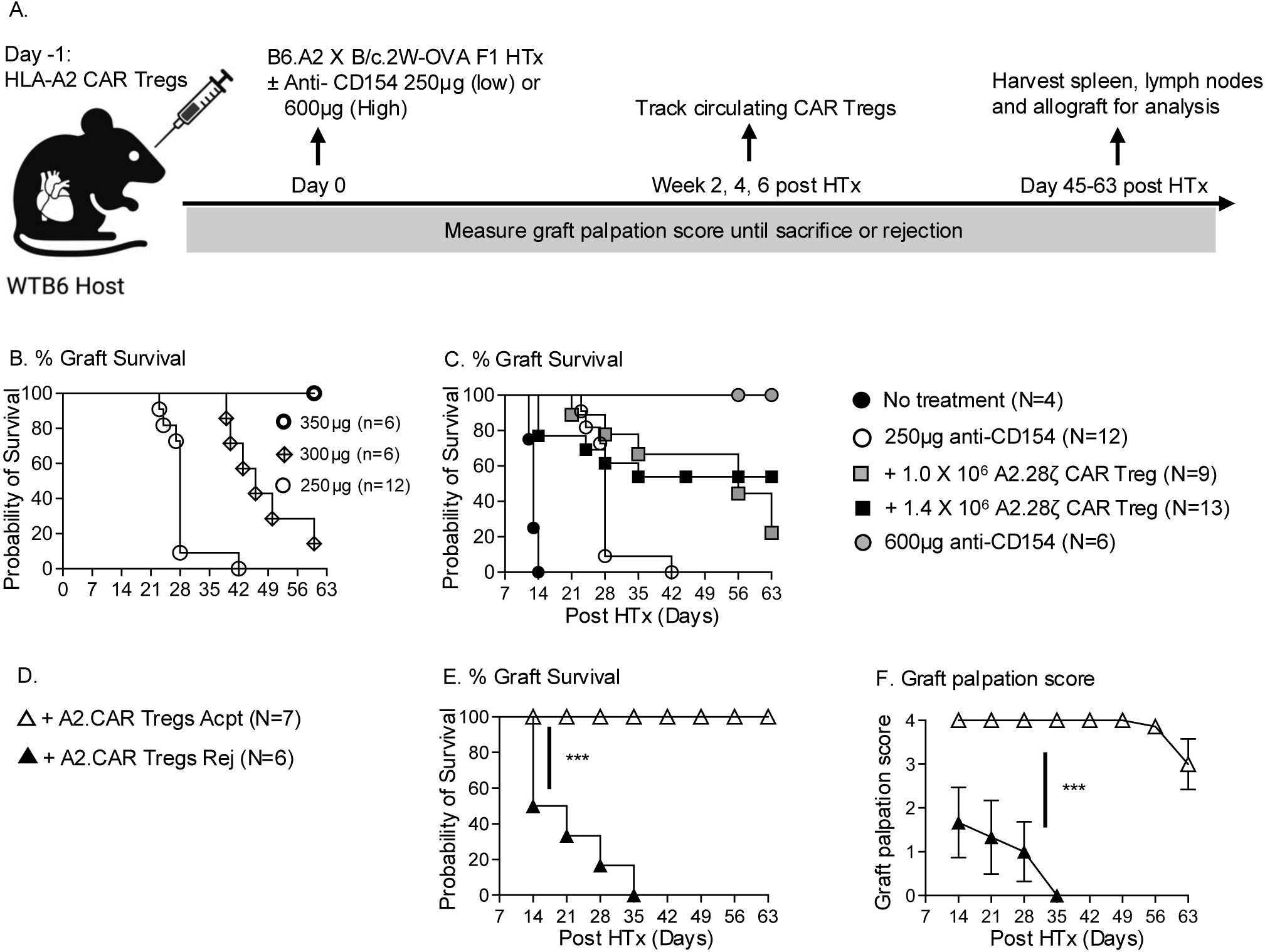
A2.CAR Tregs synergize with low dose anti-CD154 to prolong the survival of haplo-mismatched heart grafts. **(A)** Schematic of the experimental model. **(B)** Graft survival curve in recipients of A2.2W-OVA.F1 heart transplant (HTx) with three different doses of anti-CD154. **(C)** Graft survival of B6 recipients of A2.2W-OVA.F1 HTx treated with low anti-CD154 (250µg) without or with CAR Tregs, or high anti-CD154 (600µg). **(D)** Symbols for E-F. Graft survival **(E)** and palpation scores **(F)** in recipients of 1.4×10^6^ A2.CAR Tregs + low anti-CD154 with rejected (Rej) or accepted (Acpt) grafts. Log-rank (Mantel-Cox) test for statistical significance. Each symbol represents one mouse. Data are presented as mean ± SEM and statistical significance assessed by Mann-Whitney Test. *P <0.05; **P <0.01; ***P <0.001.

Recipients of 1.0×10^6^ A2.CAR Tregs + anti-CD154 had significantly prolonged median survival of 56 days as did a higher dose of 1.4×10^6^ A2.CAR Tregs (henceforth referred to as +A2.CAR Treg) (Figure 1C). However, there was considerable heterogeneity in graft survival (Figure 1D-1F), with 23% (n=3/13) rapidly rejecting their grafts by day 14, and another 23% (n=3/13) by day 35 post-HTx. The remaining 53% of recipients (n=7/13) maintained a graft palpation score of >3+ until the day of sacrifice (day 45-63 post-HTx). Thus, 1.4×10^6^ A2.CAR Tregs recipients were grouped into rejecting (Rej) and accepting (Acpt) groups for subsequent characterization of A2.CAR Tregs and endogenous donor-specific T and B cell responses.

### Rejection in A2.CAR Treg recipients is associated with reduced FoxP3^pos^:FoxP3^neg^ A2.CAR T cell ratios within heart allografts

Heterogeneity in graft survival in A2.CAR Treg recipients prompted us to track circulating CD4^+^Thy1.1^+^ A2.CAR Tregs at weeks 2-6 post-HTx (Supplementary Figure 1D). At all timepoints, a surprising ∼10-fold higher number of CD4^+^Thy1.1^+^ A2.CAR Tregs was detected in the blood of Rej compared to Acpt recipients (Figure 2A), although the percentage of cells expressing FoxP3 in Rej versus Acpt recipients was not significantly different across all three time points (Figure 2B). Consequently, both FoxP3^pos^ and FoxP3^neg^ circulating A2.CAR T cells were present in higher numbers in Rej compared to Acpt recipients (Figure 2C, D). This suggests that an early expansion of circulating A2.CAR Tregs was unexpectedly associated with poor allograft survival.

**Figure 2:**
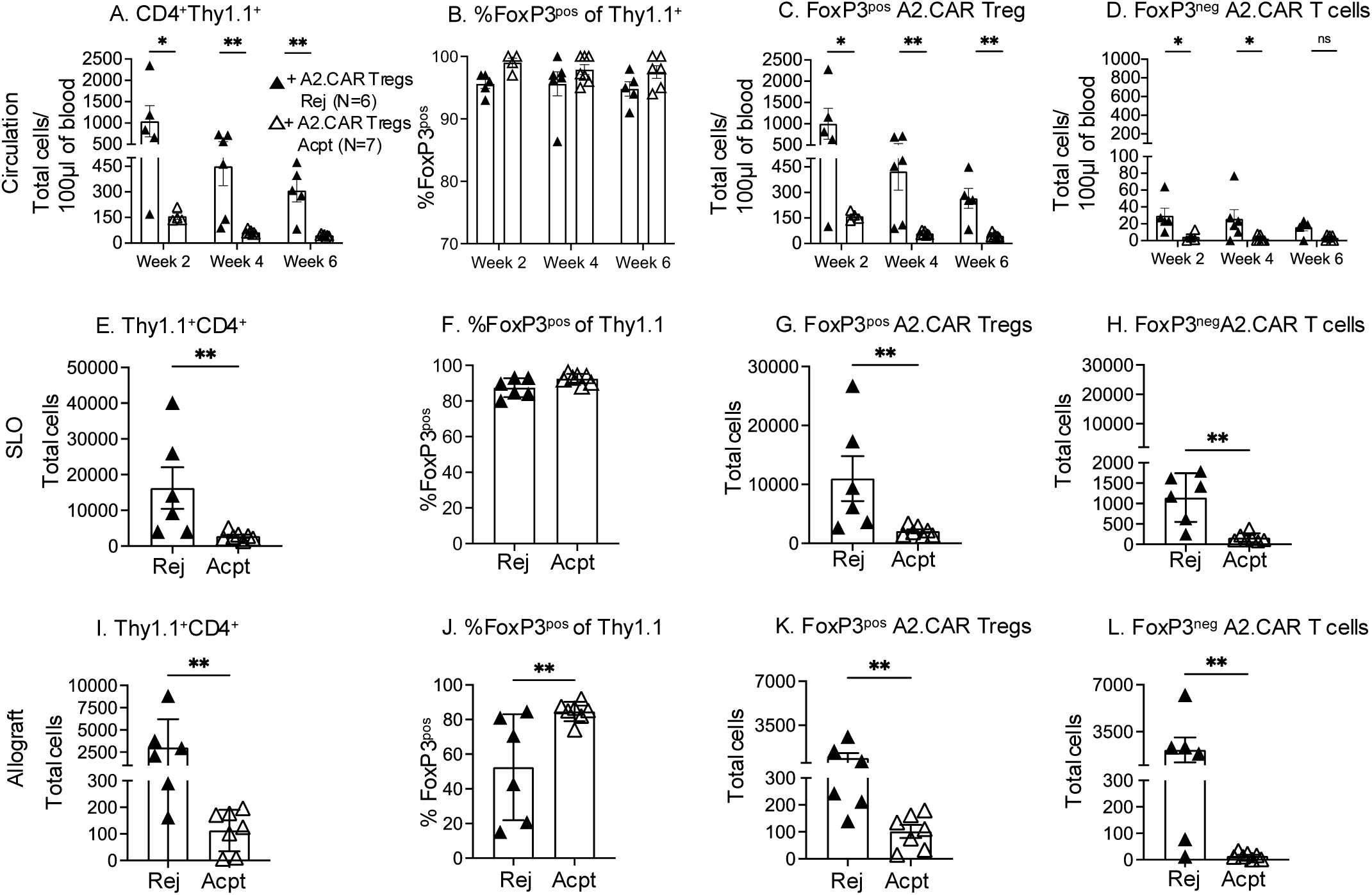
Recipients of A2.CAR Tregs and low-dose anti-CD154 with accepted allografts have low accumulation of CD4^+^Thy1.1^+^ cells and high ratios of FoxP3^pos^: FoxP3^neg^ A2.CAR T cells. (A-D) The absolute numbers and proportions of circulating Thy1.1+ A2.CAR Tregs in A2.CAR Treg recipients with rejected (Rej; filled triangles) or accepted (Acpt; open triangles) was quantified at the indicated weeks. **(A)** Number of CD4^+^Thy1.1^+^ cells per 100µl blood. **(B)** %FoxP3^pos^ of Thy1.1^+^ cells. **(C, D)** Number of FoxP3^pos^ or FoxP3^neg^ A2.CAR T cells per 100µl blood. **(E-L)** At the experimental endpoint (POD 45-63) SLO and allografts were harvested, and Thy1.1^+^ A2.CAR Tregs quantified. **(E, I)** Total number of CD4**^+^**Thy1.1**^+^** T cells from SLOs and allografts from Rej (n=6) or Acpt (n=7) recipients. **(F, J)** %FoxP3^pos^ of Thy1.1**^+^** cells. Number of FoxP3^pos^ **(G, K)** or FoxP3^neg^ **(H, L)** A2.CAR T cells recovered from SLO and allograft. Each symbol represents one mouse. Data are presented as mean ± SEM, and statistical significance assessed by Mann-Whitney Test. **P <0.01. SLO: Pooled lymph nodes and spleen. Total number of CD4**^+^**Thy1.1**^+^** cells from SLO normalized to 3×10^6^ events. Total number of CD4**^+^**Thy1.1**^+^** cells from allografts normalized to 1×10^6^ events.

To verify the association between expanded numbers of A2.CAR Tregs and poor allograft survival, we quantified FoxP3^pos^ and FoxP3^neg^ of CD4^+^Thy.1^+^ cells from the spleen and lymph nodes (SLO) as well as allografts at the time of sacrifice (day 45-63 post-HTx). Similar to blood, significantly more (6-7-fold) CD4^+^Thy1.1^+^ CAR Tregs was recovered from the SLOs of Rej versus Acpt recipients, with comparable percentages of FoxP3^pos^ of CD4^+^Thy1.1^+^ cells (87-93%). Thus, more FoxP3^pos^ and FoxP3^neg^ A2.CAR T cells were recovered from Rej vs Acpt recipients (Figure 2E-H, Supplementary figure 2A). CAR (Myc) expression was also significantly elevated on A2.CAR Tregs from the circulation and SLO of Rej versus Acpt recipients (Supplementary figure 2B-E).

The number of total CD4^+^Thy1.1^+^ cells infiltrating Rej allografts was also higher (27±28-fold) compared to Acpt recipients (Figure 2I). Notably, only 52 ± 30% of Thy1.1^+^ cells were FoxP3^pos^ in Rej compared to 85 ± 5% in Acpt allografts (Figure 2J). As a result, the recovered Foxp3^pos^ cells were only a mean 8-fold (1880±2492) higher whereas Foxp3^neg^ cells were ∼130-fold higher (3562±4368) in Rej allografts compared to Acpt (Figure 2K-L). Finally, while the overall CAR (Myc) expression by graft infiltrating CD4^+^Thy1.1^+^ T cells was lower compared to those in blood or SLO, CAR expression was significantly higher from Rej versus Acpt allografts (Supplementary Figure 2F-G).

### FoxP3^pos^ and FoxP3^neg^ A2.CAR T cells acquire distinct activation phenotypes in rejection and acceptance in the SLO

The differential expansion of Foxp3 positive versus negative A2.CAR T cells in Rej versus Acpt recipients prompted a phenotypic analysis of markers associated with Treg function or T cell dysfunction, namely FR4, CD73, PD1, SLAMF6, LAG3 and TIGIT. Uniform Manifold Approximation and Projection (UMAP) was used to visualize the phenotype at single cell resolution (Figure 3A-B) revealing that Foxp3^pos^ and FoxP3^neg^ A2.CAR T cells from Acpt versus Rej recipients were phenotypically distinct (Figure 3C-D). Specifically, Foxp3^pos^ A2.CAR Tregs from Acpt recipients had higher expression of CD44, FR4, and CD73, but comparable levels of PD-1 and SLAMF6 (Figure 3E). Thus, A2.CAR T cells in Rej recipients exhibited significantly greater expansion and acquired a phenotype suggestive of reduced regulatory function. FoxP3^neg^ A2.CAR T cells from Acpt versus Rej recipients also expressed higher FR4, but lower PD-1 and SLAMF6 (Figure 3F).

**Figure 3:**
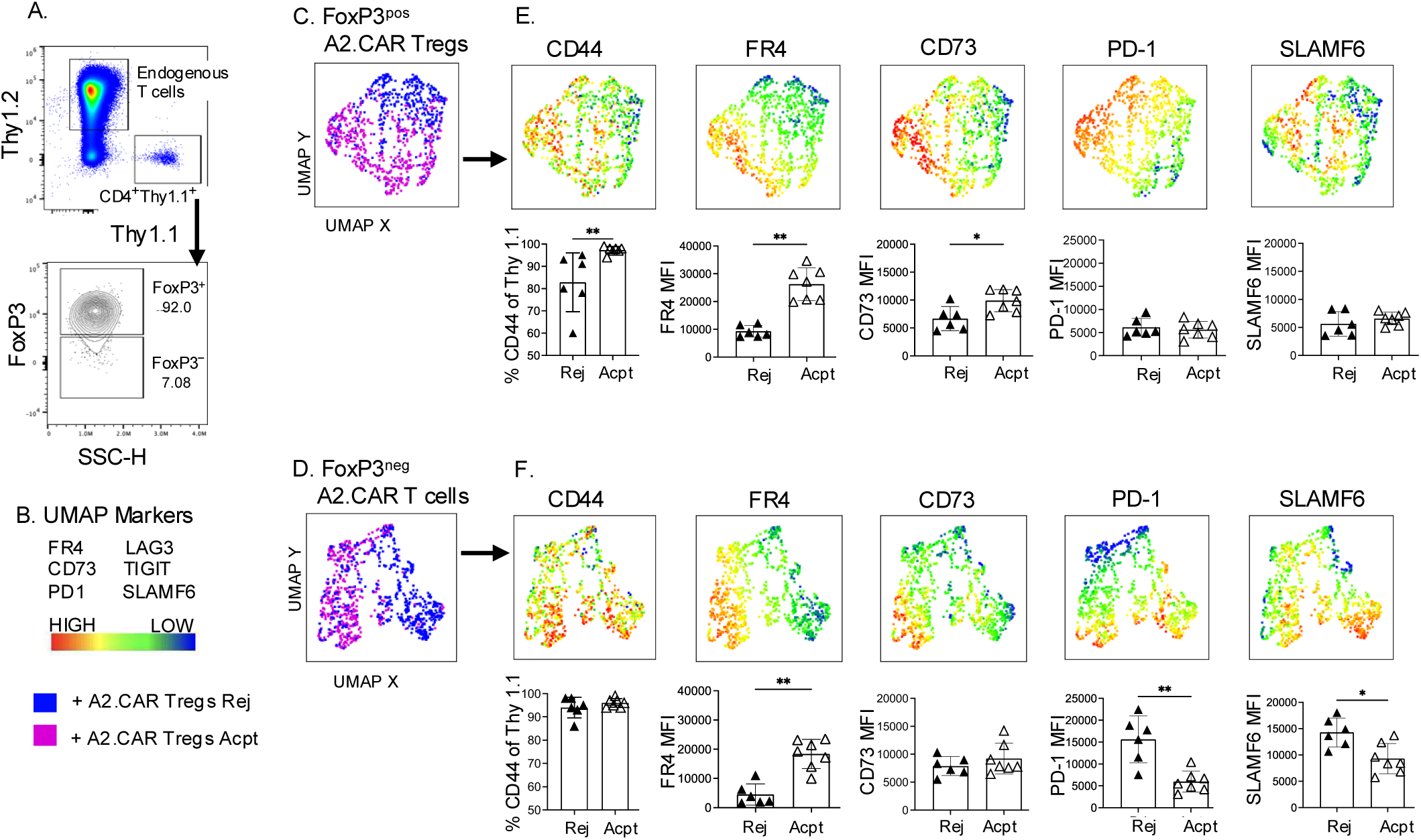
FoxP3^pos^ and FoxP3^neg^ A2.CAR T cells in the SLO are phenotypically distinct in recipients with rejecting (Rej) or accepting (Acpt) allografts. **(A)** Gating strategy to define FoxP3^pos^ and FoxP3^neg^ cells within CD4^+^Thy1.1^+^ A2.CAR T cells from SLO (POD 45-63). **(B)** UMAP markers and experimental groups. **(C, D)** UMAP plots demonstrating phenotypic differences in **(E)** FoxP3^pos^ and **(F)** FoxP3^neg^A2.CAR T cells from Rej and Acpt recipients, based on expression of FR4, CD73, PD1, SLAMF6, LAG3, and TIGIT. UMAP with heatmap and bar plots showing relative expression of indicated markers based on normalized median fluorescence intensity (MFI). Each symbol in the bar plots represents one mouse. Data are presented as mean ± SEM and statistical significance determined by Mann Whitney Test (*). *P <0.05; **P <0.01.

### A2.CAR Tregs promote endogenous Treg expansion and suppress donor-reactive Tconv and CD8^+^ T cells

In our experimental model, donor grafts expressed A2, BALB/c alloantigens, as well the model antigen 2W-OVA. The latter transgene enabled tracking of endogenous polyclonal 2W- and OVA-specific CD4^+^ and CD8^+^ T cells by utilizing 2W:I- A^b^ and OVA:K^b^ tetramers, respectively (Supplementary Figure 3). The experimental groups were Acpt or Rej HTx-recipients receiving A2.CAR Tregs and low anti-CD154, and control were naïve B6, B6 HTx recipients with no treatment (acute rejecting, AR), or receiving low or high dose anti-CD154 monotherapy. Overall, the total number of CD4^+^ 2W:I-A^b^ CD44^hi^ T cells recovered from the SLOs of Acpt recipients was significantly lower compared to Rej recipients (Figure 4B). Remarkably, the number of Foxp3^pos^ 2W:I-A^b^ Tregs recovered was significantly higher while the number of Foxp3^neg^ 2W:I-A^b^ Tconvs was lower in Acpt versus Rej recipients (Figure 4C-F). Consequently, the percentage of Foxp3^pos^ of 2W:I-A^b^ cells was significantly higher in Acpt versus Rej recipients (35-56% versus 5-15%). The total number of OVA:K^b^ CD8^+^ T cells was also significantly lower in Acpt recipients compared to Rej recipients (Figure 4G). These data demonstrate the ability of A2.CAR Tregs to mediate infections tolerance by reshaping host immunity towards allograft tolerance.

**Figure 4:**
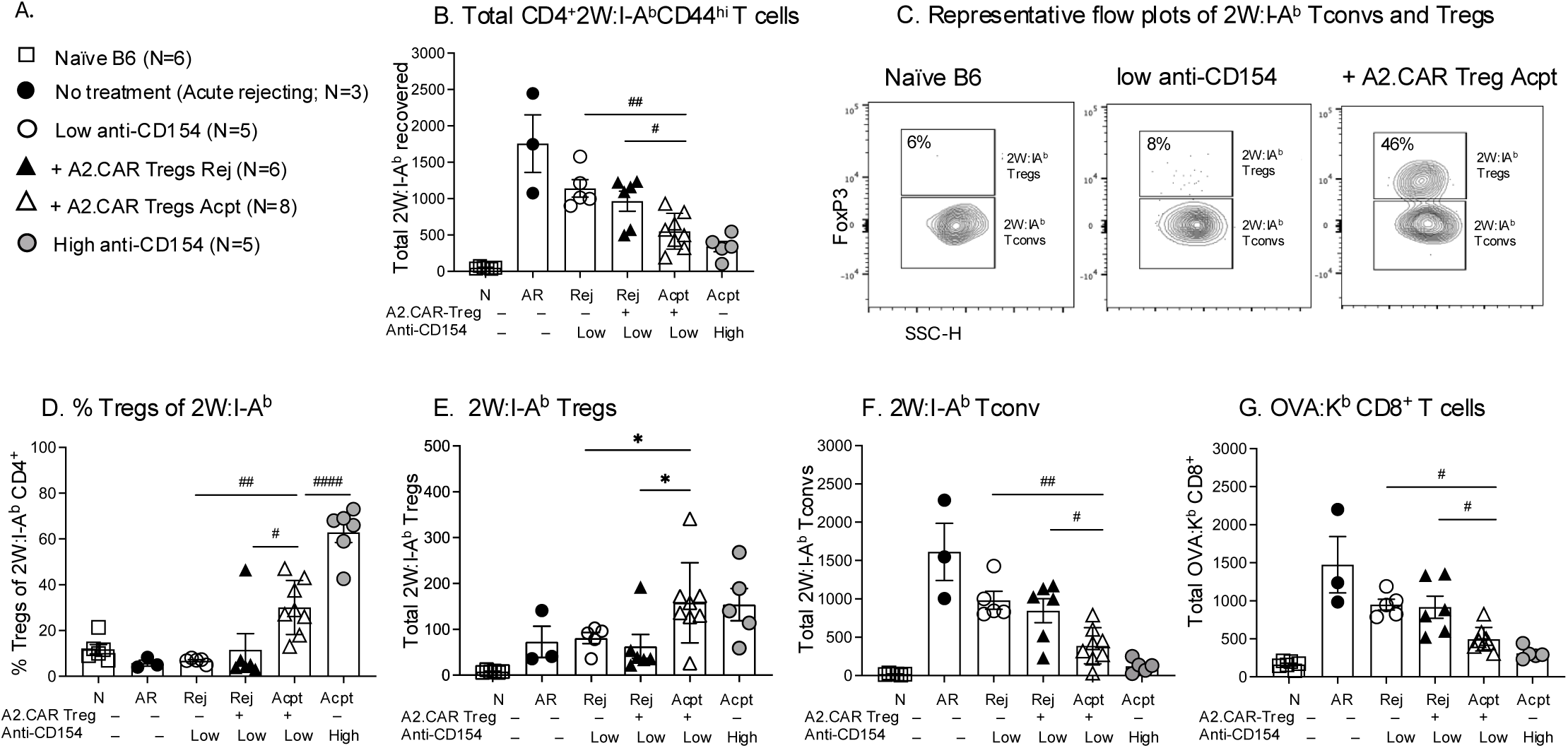
Graft acceptance in A2.CAR Tregs recipients is associated with expanded endogenous donor-specific Tregs and reduced T cell responses. **(A)** Symbols for B and D-G. **(B)** Number of CD4^+^2W:I-A^b^CD44^hi^ T cells recovered from SLOs. **(C)** Representative flow plots of FoxP3^pos^ of CD4^+^2W:I-A^b^ CD44^hi^ T cells from SLOs harvested on day 45-63 post-HTx. **(D, E)** Frequency and total number of 2W:I-A^b^CD44^hi^ Tregs, and of **(F)** 2W:I-A^b^CD44^hi^ T Tconvs and **(G)** OVA:K^b^ CD8^+^ T cells. Each symbol represents one mouse. (7 Acpt received 1.4 ×10^6^, and 1 Acpt at day 100 post-HTx received 1×10^6^ A2.CAR Tregs). Total 2W:I-A^b^ CD4^+^ T cells and OVA:K^b^ CD8^+^ T cells from SLOs normalized to 3×10^6^ events. Data are presented as mean ± SEM and statistical significance determined by 1 way ANOVA (#) and Mann Whitney Test (*). *P or ^#^P <0.05; ^##^P <0.01; ^####^P <0.0001.

Flow cytometric analysis of 2W:I-A^b^ Tregs and Tconvs, as well as OVA:K^b^ CD8^+^ from Acpt versus Rej recipients of A2.CAR Tregs + anti-CD154 revealed that they were phenotypically comparable (Supplementary figure 4A, 5A, 6A) . This panel of markers, namely FR4, CD73, PD-1 and SLAM6, was able to distinguish these T cells from control rejection (untreated or low anti-CD154) versus control acceptance (high anti-CD154; Supplementary Figures 4B, 5B, and 6B). Thus, while endogenous donor-specific CD4^+^ and CD8^+^ T cells expanded in Rej recipients, their conserved phenotype in Rej and Acpt prompted us to next assess if they were able to infiltrate the graft to participate in graft rejection.

Histology of transplanted hearts (Figure 5A-E) confirmed that lymphocyte infiltration was minimal in Acpt but abundant in Rej grafts. Flow cytometry was performed to quantify the endogenous (Thy1.1^neg^) T cells and B cells accumulating in allografts confirmed that endogenous T cell and B cell infiltration was significantly increased in Rej versus Acpt grafts (Figure 5F-I). Taken together, these data illustrate the ability of A2.CAR Tregs to synergize with anti-CD154 to mediate infectious tolerance leading to controlled endogenous donor-specific CD4⁺ and CD8⁺ T cell responses, both systemically and in the graft, and to improved allograft survival. In contrast, no infectious tolerance was observed in A2.CAR Tregs recipients that rejected their grafts.

**Figure 5:**
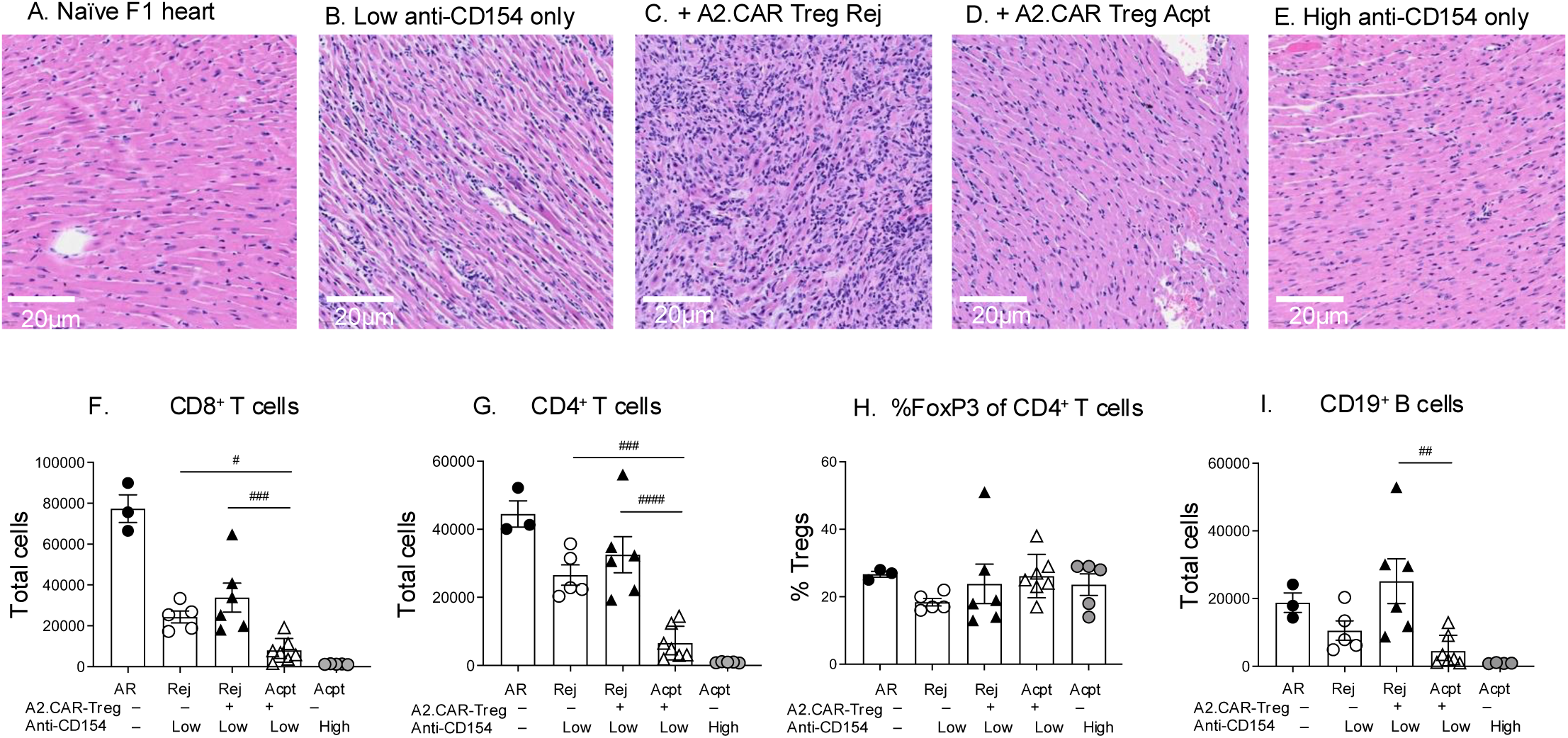
Graft acceptance in A2.CAR Treg recipients is associated with reduced accumulation of endogenous T and B cells in heart allografts. (A-E) Representative H&E-stained histological sections of naïve or transplanted heart allografts from indicated groups sacrificed on days 45-63 post-HTx. **(F-I)** Cells infiltrating the allograft were analyzed by flow cytometry. Total graft infiltrating **(F)** CD8**^+^** T cells, **(G)** CD4**^+^** T cells, **(H)** % Tregs of CD4^+^ T cells, and **(I)** CD19^+^ B cells were normalized to 1×10^6^ events. Each symbol represents each mouse. Data are presented as mean ± SEM, and statistical significance assessed by one-way-ANOVA. ^#^P <0.05; ^##^P <0.01; ^###^P <0.001.

*A2.CAR Tregs inhibit anti-A2 and anti-donor MHC B cell responses.* Prior studies reported that A2.CAR Tregs suppressed anti-A2 IgG responses in single antigen-mismatched transplant models (21), but the ability to control non-A2 IgG was unknown. We found that in Acpt recipients A2.CAR Tregs significantly suppressed DSA responses to HLA-A2 as well as to BALB/c MHC-I (K^d^, L^d^) and MHC-II (I-E^d^, I-A^d^) (Figure 6A-C). Indeed, IgG levels to all three specificities were comparable to recipients treated with high anti-CD154 or naive mice. In contrast, Rej recipients had higher levels of DSA by day 45 post-HTx, although anti-donor MHC I titers remained significantly lower compared to recipients of low anti-CD154 without A2.CAR-Tregs.

**Figure 6:**
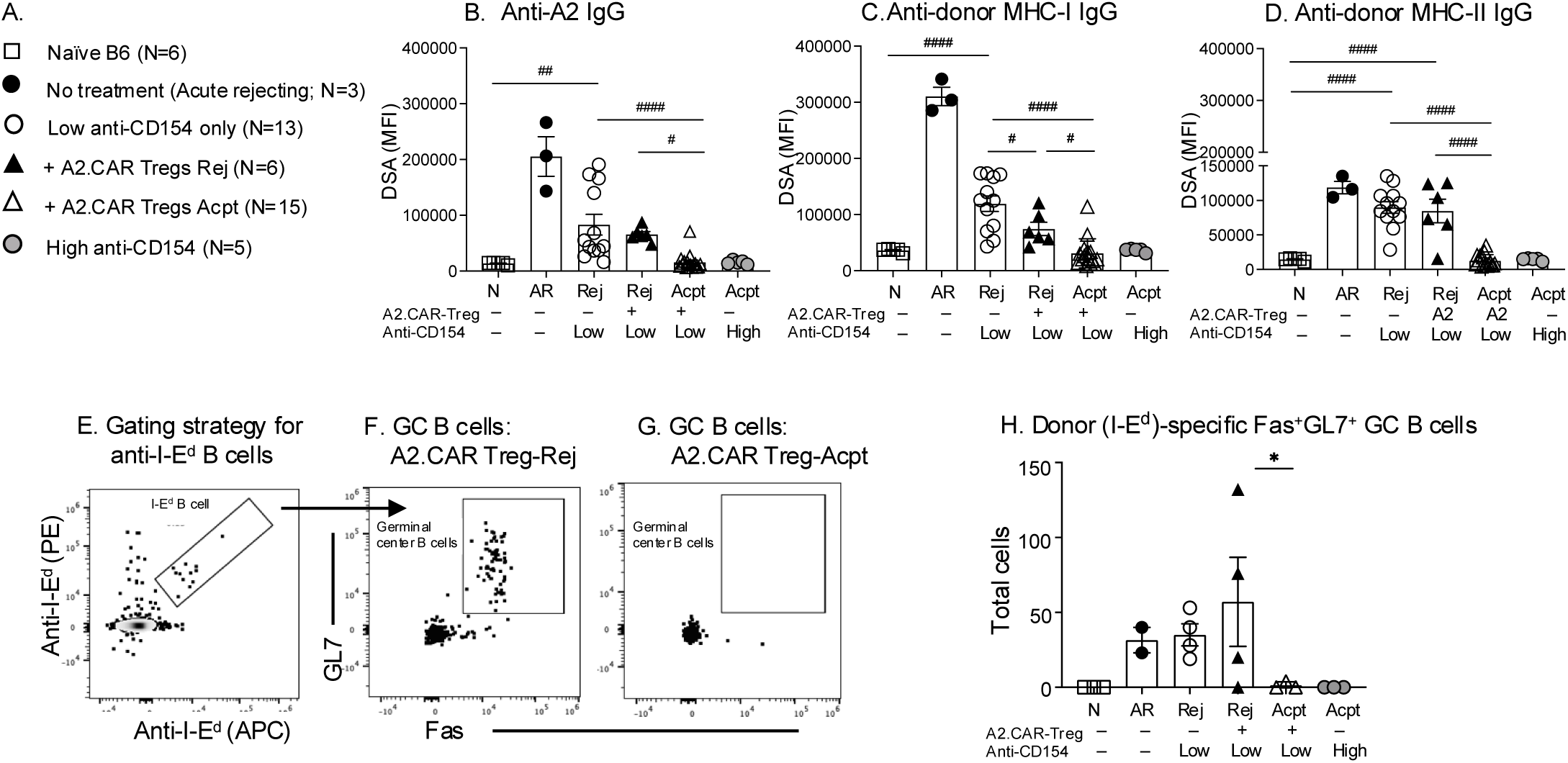
A2.CAR Tregs synergize with low anti-CD154 to suppress non-A2 donor- specific B cell responses. **(A)** Symbols for B-D and H. **(B)** Quantification of IgG specific for A2, **(C)** donor MHC Class I IgG and **(D)** donor Class II at day 45 post HTx. **(E-G)** Gating strategy to track donor MHC-II I-E^d^-specific GC B cells (Fas^+^GL7^+^) from **(F)** +A2.CAR Treg Rej or **(G)** +A2.CAR Treg Acpt. **(H)** Total GC B cells (Fas^+^GL7^+^ of I-E^d^) recovered from draining lymph nodes (normalized to 2×10^6^ events). Each symbol represents one mouse. Data are presented as mean ± SEM and statistical significance determined by 1 way ANOVA (#) and Mann Whitney Test (*). *P or ^#^P <0.05; ^##^P <0.01; ^####^P <0.0001. MFI: Median fluorescence intensity. DSA: Donor specific antibody.

Furthermore, by utilizing donor MHCII I-E^d^ tetramers, we tracked endogenous I-E^d^ specific B cells and characterized their phenotype (Figure 6D-G). Recipients with Acpt allografts had decreased germinal center B (GC: Fas^+^GL7^+^) responses compared to recipients with Rej allografts or recipients treated with low anti-CD154 only. Thus, these data show that A2.CAR Treg therapy is able to control endogenous DSA responses that are specific not only for A2, but also for donor MHC class I and II.

## Discussion

CAR Tregs are already being tested clinically, despite limited experimental evidence for how to effectively combine them with immunosuppression to promote the stable acceptance of multiple antigen-mismatched allografts in immunocompetent recipients(22). Our study provides major advances in both of these areas by providing evidence that in combination with CD154 blockade, A2.CAR Tregs extend the survival of haplo-mismatched heart allografts. Importantly, treatment with CAR Tregs specific for a single donor (A2) antigen suppressed non-A2 donor-reactive CD4, CD8 and B cell responses and promoted the expansion of FoxP3^+^ cells within the endogenous donor-specific (2W:I-A^b^) CD4^+^ T cell population. These data build upon evidence that A2.CAR Treg therapy promotes infectious tolerance in a model of islet transplantation (27) and support the conclusion that infusion of a single dose of donor-specific CAR Tregs can broadly re-shape allograft immunity to promote tolerance. Since the general consensus is that long-term allograft tolerance is mediated by endogenous Tregs with indirect specificity (28, 29), evidence that infused A2.CAR Tregs can promote the expansion of endogenous Treg with indirect specificity is a key advance. A caveat is that the A2.CAR Tregs also express endogenous TCR, so we cannot rule out the possibility that some of the effects may be mediated by endogenous TCRs with indirect alloreactivity.

It was previously reported that A2.CAR Tregs can suppress A2-specific B cells and antibody production (21), but their ability to control antibody responses to other alloantigens was unknown. Here, we showed that both A2-specific and IgG responses specific for donor MHCI and II were successfully controlled, and that the differentiation of donor MHC-specific B cells into GC B cells was inhibited compared to recipients treated with low anti-CD154 alone. Whether this control of donor-specific humoral immunity was mediated by A2.CAR Tregs preventing the generation of T follicular helper (Tfh) cells or promoting the differentiation of regulatory T follicular regulatory (Tfrs) or both will require further investigation (30). Nevertheless, the observation that infectious tolerance extends to B cell responses is important as DSA has been implicated in chronic allograft rejection in the clinic (31–33).

It is notable that the beneficial effects of anti-CD154 and A2.CAR Treg therapy only occurred in approximately half of the recipients. This finding aligns with other studies where the efficacy of adoptively transferred Tregs in models of islet, heart, and skin transplants was also heterogenous and graft survival rates did not exceed 80% at day 100 post-transplantation (29, 34, 35). Investigations into mice where allograft survival was not prolonged showed no evidence for infectious tolerance. Rather, as early as 2 weeks post-adoptive transfer, there was a significant expansion in A2.CAR-Tregs in the blood, and SLOs of Rej compared to Acpt recipients, although the percentage of cells that were FoxP3^pos^ remained comparable. We also observed higher ratios of FoxP3^neg^:FoxP3^pos^ A2.CAR T cells within Rej versus Acpt grafts at the time of sacrifice (45-63 days post-HTx), thus emphasizing the need to investigate events within the allograft. This accumulation of FoxP3^neg^ A2.CAR T cells could be due to preferential expansion of FoxP3^neg^ cells present in the infused product, which contained ∼3-5% Foxp3^neg^ cells, and/or destabilization of FoxP3^pos^ into FoxP3^neg^ cells (36–38). Investigations into the origin of FoxP3^neg^ A2.CAR Tregs, and why A2.CAR Tregs failed to re-shape host immunity in some mice will be important future directions.

Tregs control antigen-presenting cells (APCs) and alloreactive T and B cells through a variety of mechanisms including the expression of coinhibitory molecules, trogocytosis of stimulatory molecules on APCs, secretion of immunoregulatory cytokines, depletion of IL-2, and expression of ectonucleotidases, CD73 and CD39, that catabolize ATP into immunosuppressive adenosine (39–43). Furthermore, FR4, a surface receptor for folic acid (Vitamin B9) has been implicated in mediating Treg function and is highly expressed on the antigen-responsive Tregs (44). We observed that HTx acceptance was associated with FoxP3^+pos^ A2.CAR Tregs with upregulated FR4 and CD73, raising the possibility that these represent CAR Tregs with regulatory function.

In summary, our study addresses several mechanistic gaps in knowledge and identifies new ones. A2.CAR-Tregs synergized with anti-CD154 to promote infectious tolerance. However, whether this is driven by A2.CAR Tregs secreting anti-inflammatory cytokines such as IL-10, IL-35, TGF-β that act on T cell, or by acting on APCs has yet to be identified. Second, we observed increased ratios of FoxP3^neg^:FoxP3^pos^ A2.CAR Tregs in rejecting graft whereas very few A2.CAR Tregs infiltrated accepted grafts. The inflammatory milieu unique to the rejecting graft that drives the recruitment A2.CAR Tregs and the increase in FoxP3^neg^:FoxP3^pos^ ratios remains to be elucidated. Third, it is unclear if B cell suppression is due to reduced Tfh help or direct A2.CAR Treg binding to cross-dressed HLA-A2 on B cells (45), and warrants additional study. Finally, our study supports the clinical testing of CAR Treg therapy in combination with CD154 pathway blockade. Heterogeneity in efficacy underscores the need for additional mechanistic studies to guide iterative improvement of the CAR Treg product to maximize its potency and consistency.

## Methods

### Mice

Heart transplant recipients were eight- to twelve-week-old female C57Bl/6 (B6, H- 2^b^) mice purchased from Harlan Sprague Dawley (Indianapolis, IN). Heart donor (A2.2W- OVA.F1) mice were generated by crossing 2W-OVA.BALB/c males(46) (2W-OVA.B6 mice were a gift from James J Moon, Massachusetts General Hospital, Harvard Medical School, Charlestown, Massachusetts, USA) with B6.HLA-A*0201 females (Taconic Biosciences, Germantown, NY) to generate 2W-OVA.BALB/c mice. B6-Foxp3^gfp^ Thy1.1 mice used for generating Tregs, were bred in-house(18, 21). All animal experiments were approved by the Institutional Animal Care and Use Committee at the University of Chicago or University of British Columbia and adhered to the standards of the NIH Guide for the Care and Use of Laboratory Animals or Canadian Council of Animal Care.

### Generation of A2.CAR Tregs

A2.CAR Tregs were generated as described previously(9, 18). Briefly, lymph nodes and spleen from female B6-Foxp3^gfp^ Thy1.1 mice were collected and CD4^+^ T cells were isolated by negative selection (STEMCELL Technologies). Tregs were sorted as live CD4^+^CD8^–^Thy1.1^+^Foxp3^gfp+^ or CD4^+^CD8^–^Thy1.1^+^CD25^+^CD62L^+^ cells using a Moflo Astrios cell sorter (Beckman Coulter), stimulated with anti-CD3/CD28 Dynabeads (Thermo Fisher Scientific), expanded with recombinant human IL-2 (1,000 U/mL; Proleukin) in the presence of rapamycin (50 nmol/L; Sigma-Aldrich). After 2 days, cells were transduced with a retrovirus encoding an A2-specific, CD28-containing second-generation A2.CAR (Supplementary Figure 1A). Dynabeads were removed on day 7 and CAR expression and Treg purity were determined (Supplementary Figure 1B- C). Cells were cryopreserved prior to injection as a cell therapy.

### Adoptive transfer of A2.CAR Tregs and heterotopic heart transplantation

A2.CAR Tregs were administered on the day prior to HTx at the indicated doses via the intravenous (i.v) route. Heterotopic HTx were performed as described(47), by anastomosing female donor A2.2W-OVA F1 hearts to the inferior vena cava and aorta in the peritoneal cavity of female B6 recipients. Anti-CD154 (MR1, BioXCell) was injected i.v on the day of HTx at the indicated doses. Allograft survival was monitored twice weekly starting at post-operative day (POD) 7 by transabdominal graft palpation and scored from a scale of zero to four. Allograft rejection was defined as the last day of palpable heartbeat.

### Harvesting of allograft, secondary lymphoid organs, and tissue processing for flow cytometry

Mice were euthanized at the indicated times post-HTx and heart grafts were harvested and perfused with cold PBS, disintegrated and digested by incubating for 20 min at 37°C with 2mg/ml collagenase type IV (Sigma Aldrich). Secondary lymphoid organs (SLO) comprising spleens and pooled lymph nodes (brachial, inguinal, axillary and mediastinal) were harvested, and single cell suspensions were obtained from passing through a 40µm cell strainer (Corning Inc., USA). Red blood cells were lysed via 2–3- minute incubation with ammonium chloride-potassium (ACK) lysis buffer (Quality Biological). T cells were enriched with a Pan-T cell isolation kit II (Miltenyi Biotech) and passed through LS columns on a QuadroMACS separator (Miltenyi Biotech), followed by elution with MACS buffer (2%FBS + 2mM EDTA).

### Tetramer, fluorescent antibodies and reagents

After T cell enrichment, cells were stained with Fixable Aqua Live/Dead staining (Thermo Fisher Scientific OR Invitrogen), then Fc blocked (anti-CD16/32, S17011E /Biolegend). Tetramer staining was carried out for 40- 45 min at 4°C; APC and PE-conjugated tetramers (NIH tetramer core facility). APC- and PE-conjugated 2W (EAWGALANWAVDSA):I-A^b^ tetramers were used to track allo- reactive CD4^+^ and OVA (SIINFEKL):H-2K^b^ for CD8^+^ T cells. pCons-CDR1 (FIEWNKLRFRQGLEW):I-E^d^ tetramers (PE-conjugated) and H2-K^b^ (SIINFEKL) decoy (AF647-PE-conjugated) tetramers were used to identify donor MHC-II-reactive B cells as previously described(48, 49).

Antibodies used to define the phenotype of T cells included: <u>Invitrogen:</u> CD44- BUV737 (IM7), CD8α-PE (53-6.7); <u>Abcam:</u> c-Myc- AF488 (Y69); <u>Biolegend:</u> IgG2a-PeCy7 (RMG2a-62), IgD-BV605 (11-26c.2a), T and B cell activation marker-FITC (GL7), CD73-BV605 (TY/11.8), PD1-APC-Cy7 (RMP1-30), TIGIT-PECy7 (1G9), LAG3-BV785 (C9B7W), CD62L APC (MEL-14); <u>BD Biosciences:</u> CD19-BV421 (1D3), CD95- BUV737 (Jo2), CD90.2-BUV395 (53-2.1), CD4-BV510/BV786 (GK1.5), CD4-BUV496 (RMA.5), Thy1.1- BUV496 (HIS51), CD8-BUV805 (53-6.7), FR4- BV421 (12A5), FoxP3-AlexaFluor532/PECy7 (FJK-16s), CD25 BV421 (PC61); and <u>Ablab UBC:</u> c-Myc-AF647/AF488 (9E10). Antibodies used to exclude non-T or non-B cells were CD11c- BUV661 (N418), F4/80-BUV661 1 (T45-2342), NK1.1-2 BUV661 (PK136), TER119- BUV661 (TER-119) (Biolegend).

### Multiplex bead assay to measure DSA

Multiplex streptavidin (SA) beads (PAK-5067-10K; Spherotech) were coated with biotinylated MHC-I (K^d^, L^d^), MHC-II (I-E^d^, I-A^d^) and HLA- A*0201 monomers (NIH Tetramer facility). Sera were incubated with pooled MHC I-, MHC II-, and HLA-A2^-^coated beads at 4°C for 1 hour. Beads were washed and then stained with FITC-conjugated anti-mouse IgG (1070-09; Southern Biotech). The mean fluorescence intensity (MFI) of DSA binding to the multiplex beads was determined by flow cytometry (Novocyte Quanteon, Agilent) (48).

### Statistics

Statistical significance analyses were performed using GraphPad Prism. Graft-survival significance was assessed using Kaplan-Meier/Mantel-Cox log-rank tests. Statistical differences between experimental groups were determined by 1-way-ANOVA or Mann-Whitney unpaired t test. p values ≤ 0.05 were considered statistically significant.

### Study approval

All animal experiments were approved by the Institutional Animal Care and Use Committee at the University of Chicago, and adhered to the standard of NIH *Guide for the Care and Use of Laboratory Animals* (National Academies Press, 2011).

## Author Contribution

SSD designed the study, conducted the experiments, analyzed data and wrote the manuscript. IRS, MS, MM generated A2 CAR Tregs. DY performed the heart transplantation. IS and GH provided help with data analysis. MLA provided critical feedback to the study design. MKL and ASC designed the study and wrote the manuscript. All authors edited and provided feedback on the manuscript.

## Acknowledgment

We acknowledge Qiang Wang for her assistance in the in vivo experiments. We thank the Flow Cytometry Core (RRID: SCR_017760), The Human Tissue Resource Center and Animal Resources Center at the University of Chicago for their assistance. MHC tetramers were provided by the NIH Tetramer Core Facility (contract HHSN272201300006C). This work was supported in part by grants (R01AI142747, P01AI097113) from the National Institute of Allergy and Infectious Diseases, NIH and the Canadian Institutes of Health Research (FDN-154304 and PJT-189977). SSD was funded by American Heart Association post-doctoral fellowship grant (24POST1191785). IR-S is supported by salary awards from the Canadian Institutes for Health Research and Michael Smith Health Research BC. MKL receives a salary award from the BC Children’s Hospital Research Institute and is a Canada Research Chair in Engineered Immune Tolerance.

## Data availability statement

Data are available from the corresponding author upon request.

