## Supplementary Figures for "CAR Treg synergy with anti-CD154 mediates infectious tolerance to dictate heart transplant outcomes"

### Slide 1
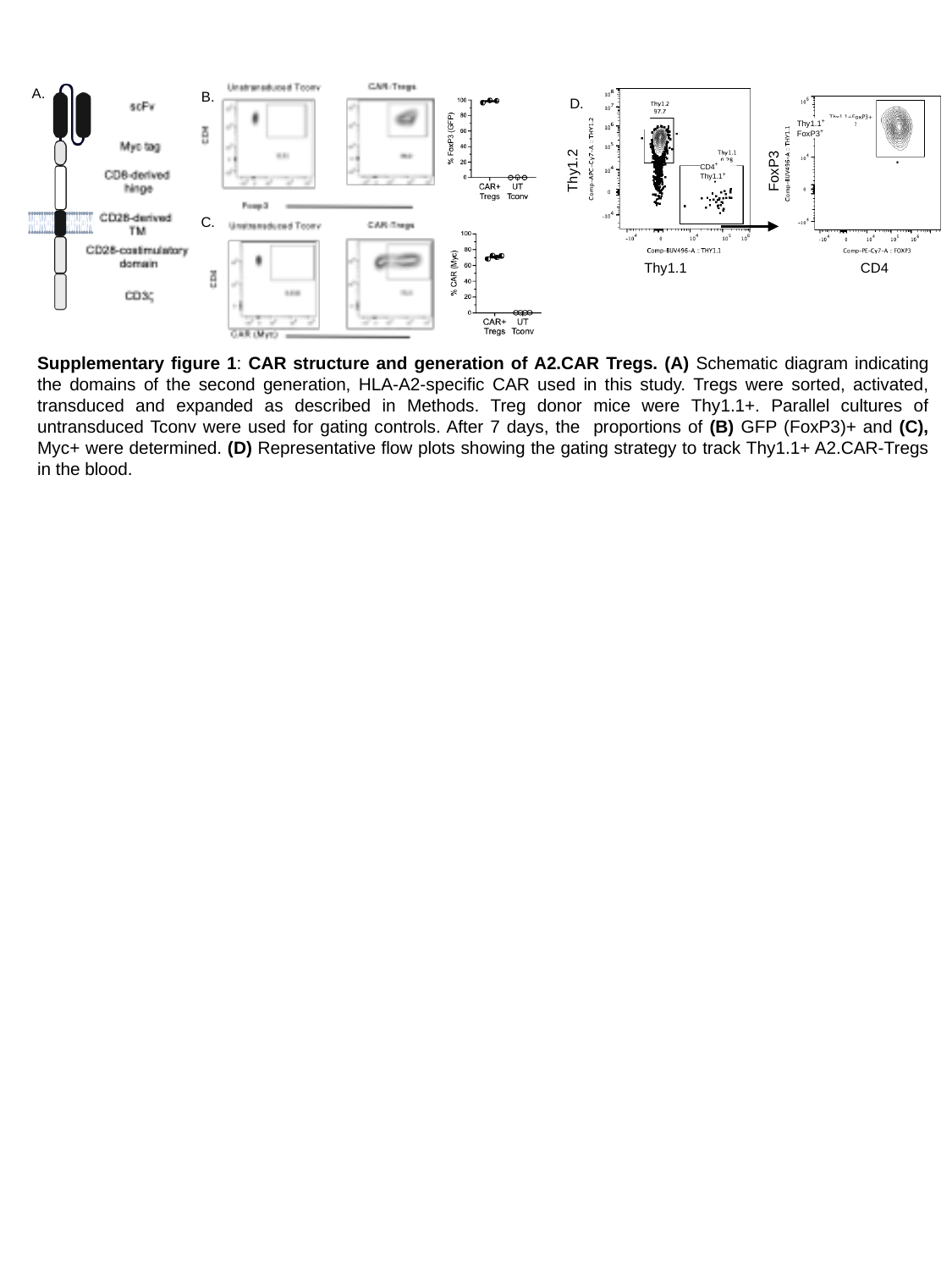

A.
B.
D.
 Thy1.1^+^
 FoxP3^+^
Thy1.2
FoxP3
CD4^+^
Thy1.1^+^
C.
Thy1.1
CD4
Supplementary figure 1: CAR structure and generation of A2.CAR Tregs. (A) Schematic diagram indicating the domains of the second generation, HLA-A2-specific CAR used in this study. Tregs were sorted, activated, transduced and expanded as described in Methods. Treg donor mice were Thy1.1+. Parallel cultures of untransduced Tconv were used for gating controls. After 7 days, the proportions of (B) GFP (FoxP3)+ and (C), Myc+ were determined. (D) Representative flow plots showing the gating strategy to track Thy1.1+ A2.CAR-Tregs in the blood.

### Slide 2
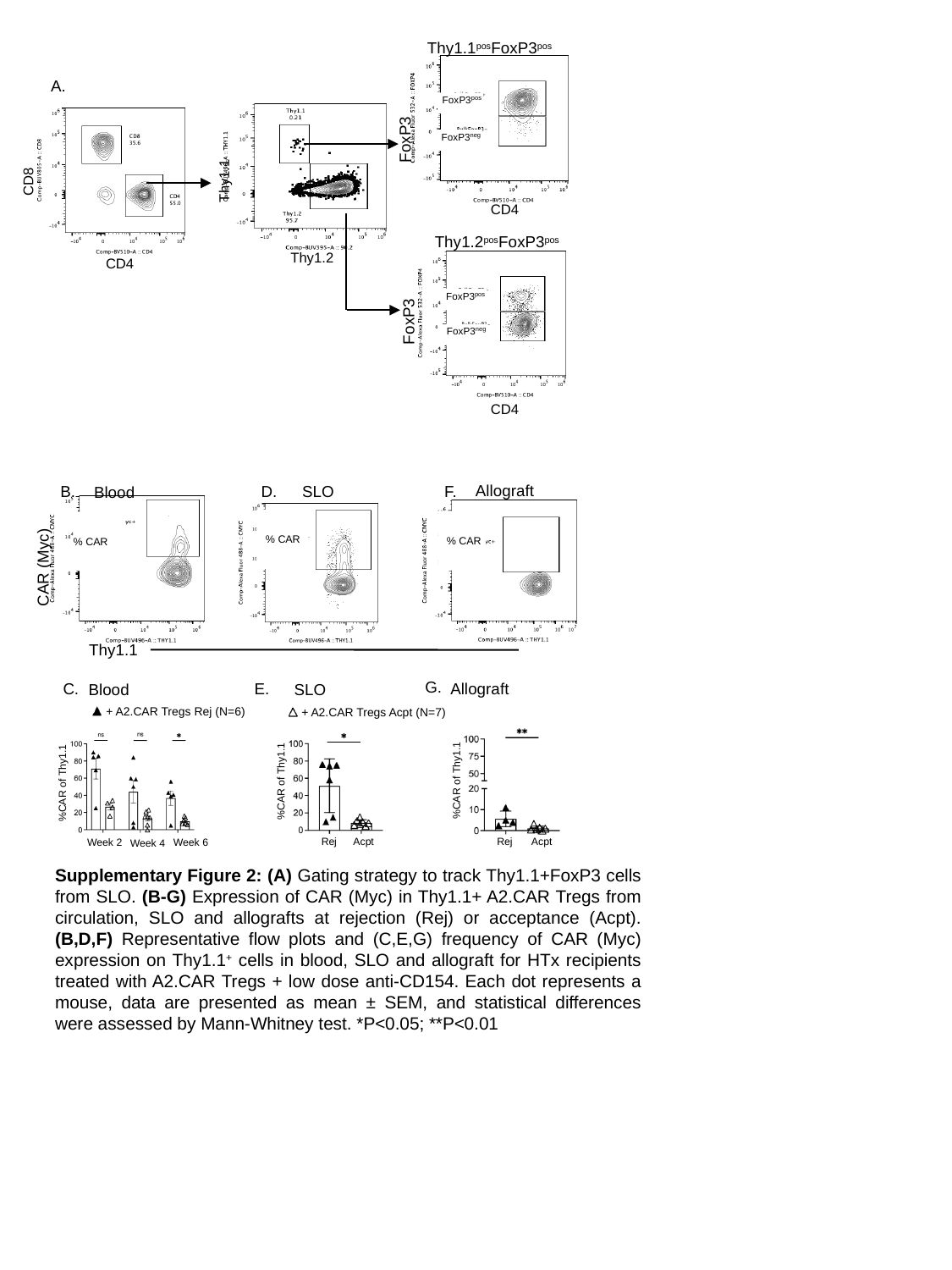

Thy1.1^pos^FoxP3^pos^
A.
 FoxP3^pos^
FoxP3
 FoxP3^neg^
Thy1.1
CD8
CD4
Thy1.2^pos^FoxP3^pos^
Thy1.2
CD4
 FoxP3^pos^
FoxP3
 FoxP3^neg^
CD4
Allograft
F.
B.
SLO
D.
Blood
 % CAR
 % CAR
% CAR
CAR (Myc)
Thy1.1
G.
C.
Allograft
E.
Blood
SLO
+ A2.CAR Tregs Rej (N=6)
+ A2.CAR Tregs Acpt (N=7)
%CAR of Thy1.1
%CAR of Thy1.1
%CAR of Thy1.1
Rej
Acpt
Rej
Acpt
Week 6
Week 2
Week 4
Supplementary Figure 2: (A) Gating strategy to track Thy1.1+FoxP3 cells from SLO. (B-G) Expression of CAR (Myc) in Thy1.1+ A2.CAR Tregs from circulation, SLO and allografts at rejection (Rej) or acceptance (Acpt). (B,D,F) Representative flow plots and (C,E,G) frequency of CAR (Myc) expression on Thy1.1^+^ cells in blood, SLO and allograft for HTx recipients treated with A2.CAR Tregs + low dose anti-CD154. Each dot represents a mouse, data are presented as mean ± SEM, and statistical differences were assessed by Mann-Whitney test. *P<0.05; **P<0.01

### Slide 3
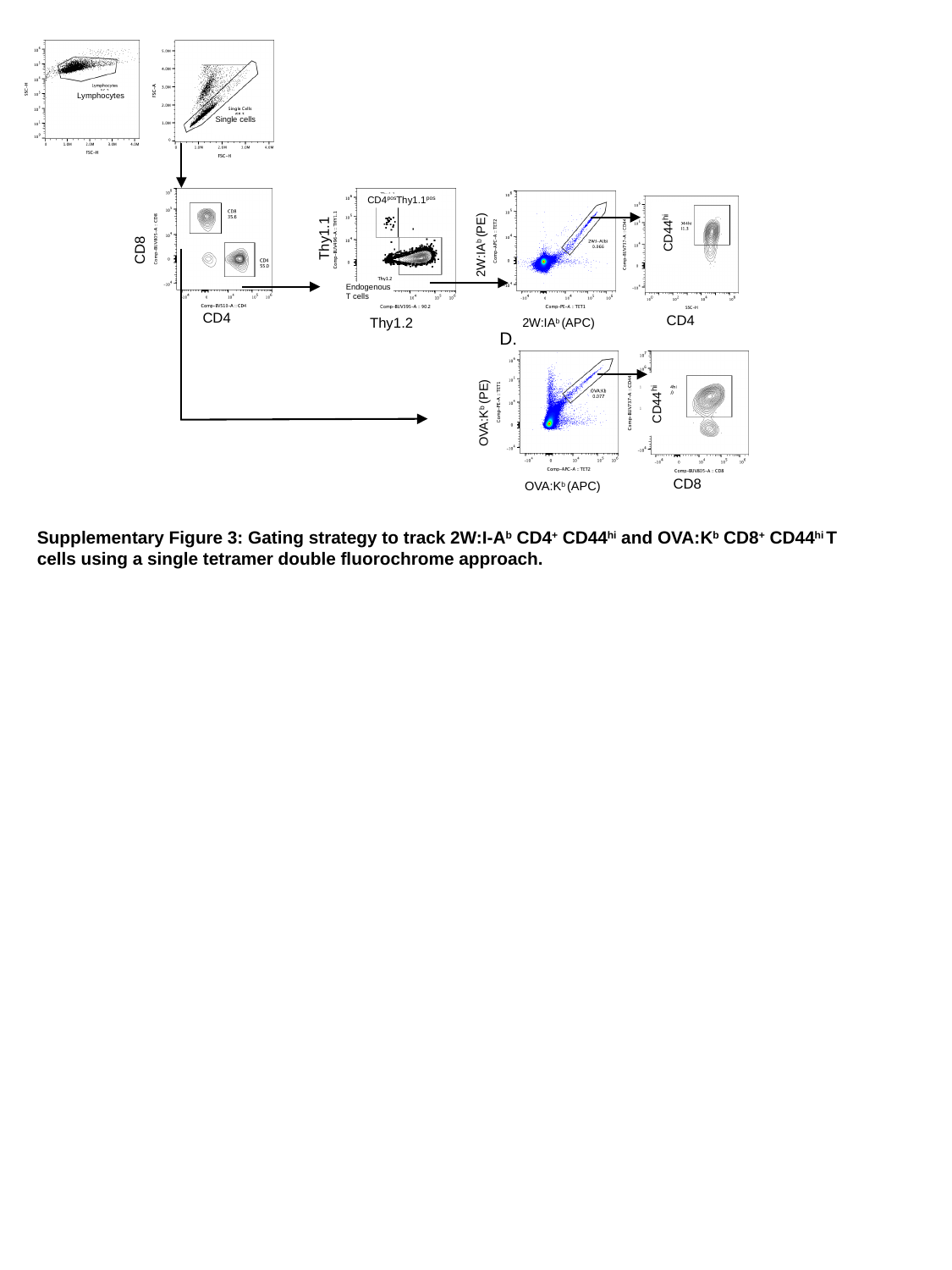

Lymphocytes
 Single cells
 CD4^pos^Thy1.1^pos^
CD44^hi^
Thy1.1
2W:IA^b^ (PE)
CD8
 Endogenous
 T cells
CD4
CD4
Thy1.2
2W:IA^b^ (APC)
D.
CD44^hi^
OVA:K^b^ (PE)
CD8
OVA:K^b^ (APC)
Supplementary Figure 3: Gating strategy to track 2W:I-A^b^ CD4^+^ CD44^hi^ and OVA:K^b^ CD8^+^ CD44^hi^ T cells using a single tetramer double fluorochrome approach.

### Slide 4
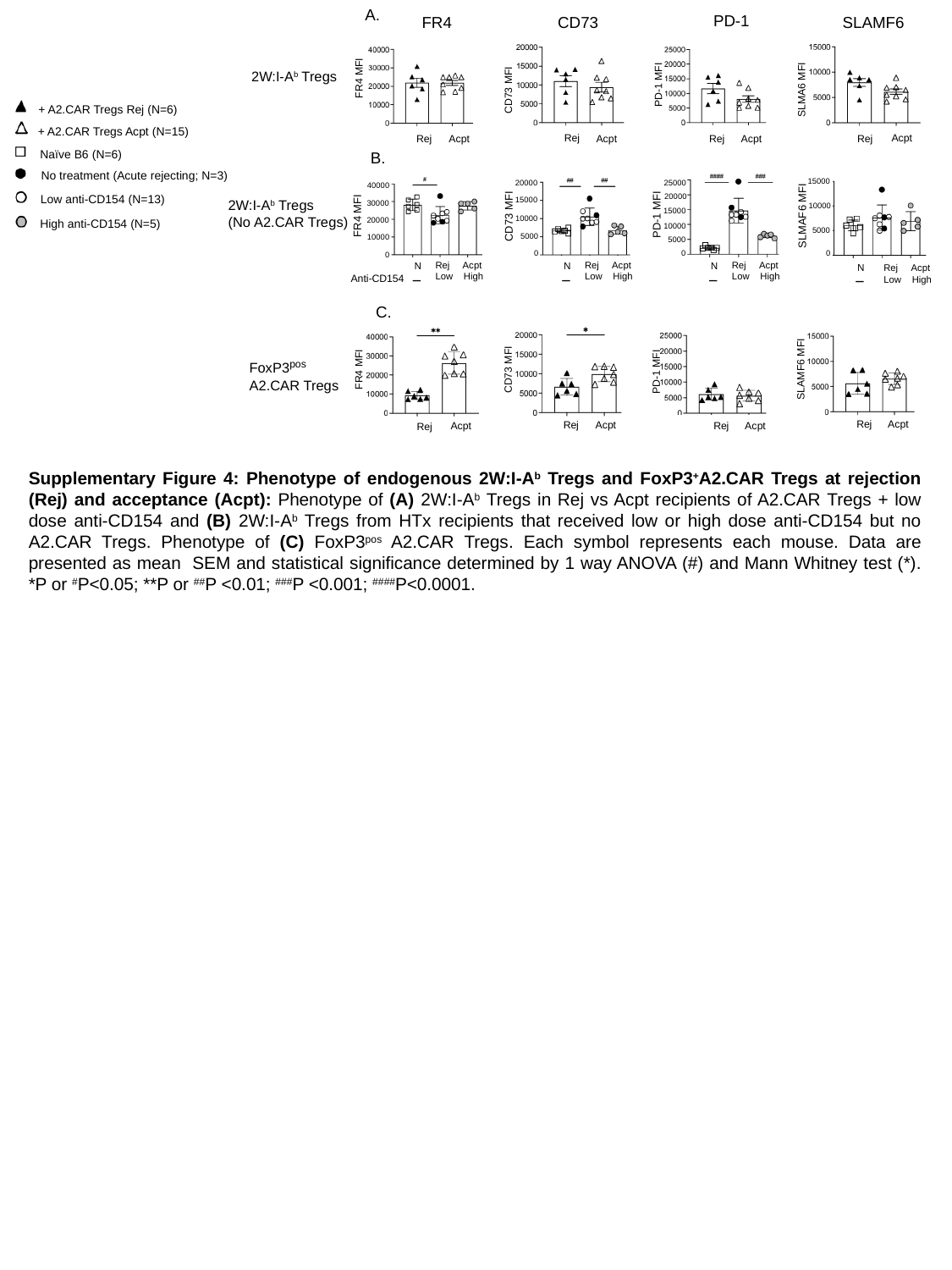

A.
PD-1
FR4
CD73
SLAMF6
2W:I-A^b^ Tregs
FR4 MFI
PD-1 MFI
CD73 MFI
SLMA6 MFI
+ A2.CAR Tregs Rej (N=6)
+ A2.CAR Tregs Acpt (N=15)
Acpt
Rej
Acpt
Rej
Rej
Acpt
Rej
Acpt
Naïve B6 (N=6)
B.
No treatment (Acute rejecting; N=3)
15000
20000
25000
40000
20000
Low anti-CD154 (N=13)
15000
2W:I-A^b^ Tregs
(No A2.CAR Tregs)
30000
10000
15000
PD-1 MFI
FR4 MFI
SLMAF6 MFI
CD73 MFI
10000
20000
High anti-CD154 (N=5)
10000
5000
5000
10000
5000
0
0
0
0
Anti-CD154
Rej Acpt Low High
Rej Acpt Low High
Rej Acpt Low High
N
N
N
_
_
_
N
_
Rej Acpt Low High
C.
FoxP3^pos^
A2.CAR Tregs
FR4 MFI
CD73 MFI
 SLAMF6 MFI
 PD-1 MFI
Rej
Acpt
Rej
Acpt
Acpt
Acpt
Rej
Rej

### Slide 5
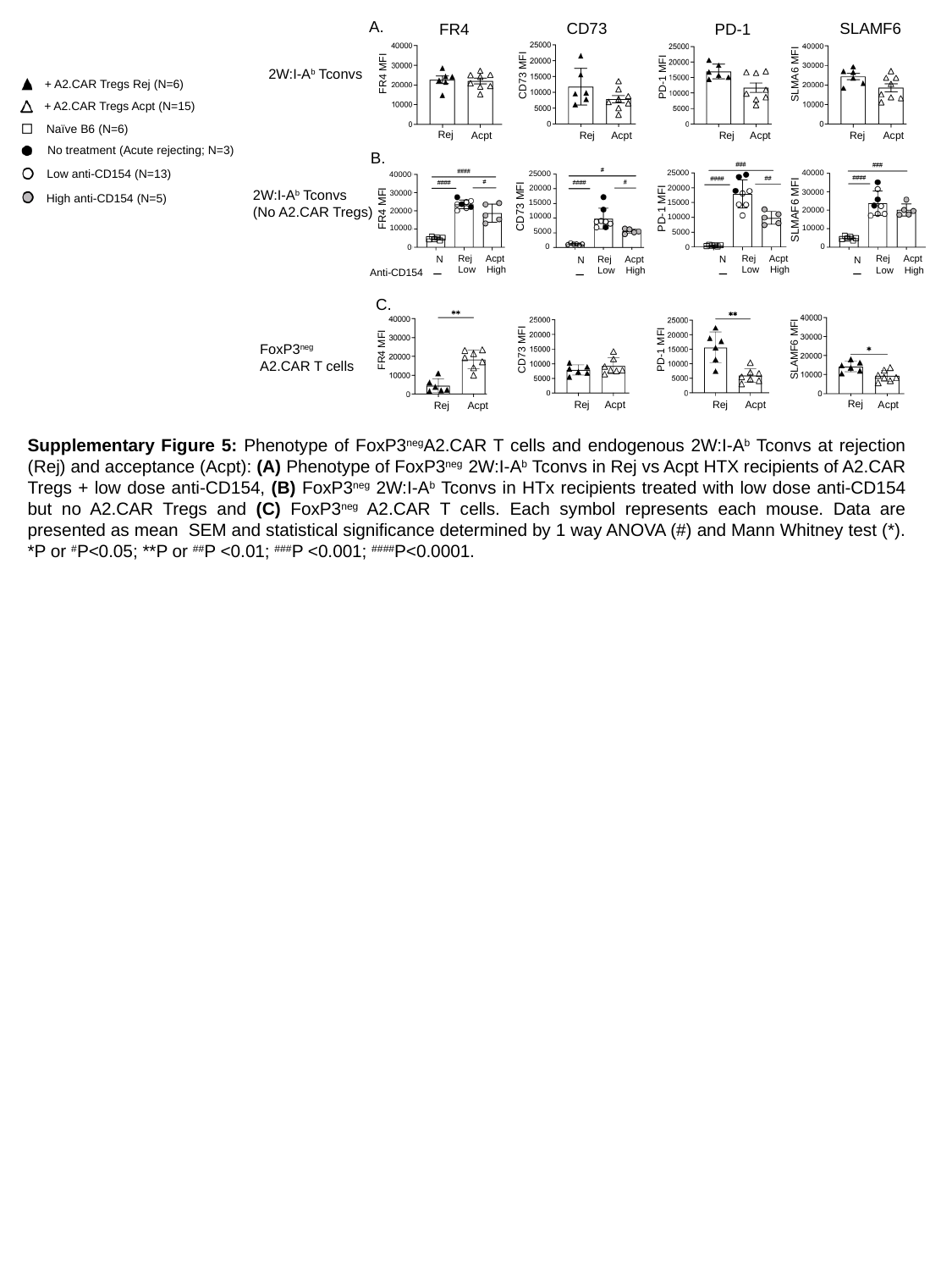

A.
SLAMF6
CD73
FR4
PD-1
2W:I-A^b^ Tconvs
FR4 MFI
SLMA6 MFI
CD73 MFI
PD-1 MFI
+ A2.CAR Tregs Rej (N=6)
+ A2.CAR Tregs Acpt (N=15)
Naïve B6 (N=6)
Rej
Acpt
Rej
Acpt
Rej
Acpt
Rej
Acpt
No treatment (Acute rejecting; N=3)
B.
Low anti-CD154 (N=13)
25000
40000
25000
40000
20000
20000
2W:I-A^b^ Tconvs
(No A2.CAR Tregs)
30000
30000
High anti-CD154 (N=5)
15000
15000
CD73 MFI
PD-1 MFI
FR4 MFI
SLMAF6 MFI
20000
20000
10000
10000
10000
10000
5000
5000
0
0
0
0
Rej Acpt Low High
Rej Acpt Low High
Rej Acpt Low High
N
Rej Acpt Low High
N
Anti-CD154
N
_
_
N
_
_
C.
FoxP3^neg^
A2.CAR T cells
CD73 MFI
FR4 MFI
 SLAMF6 MFI
 PD-1 MFI
Rej
Acpt
Rej
Acpt
Rej
Acpt
Rej
Acpt

### Slide 6
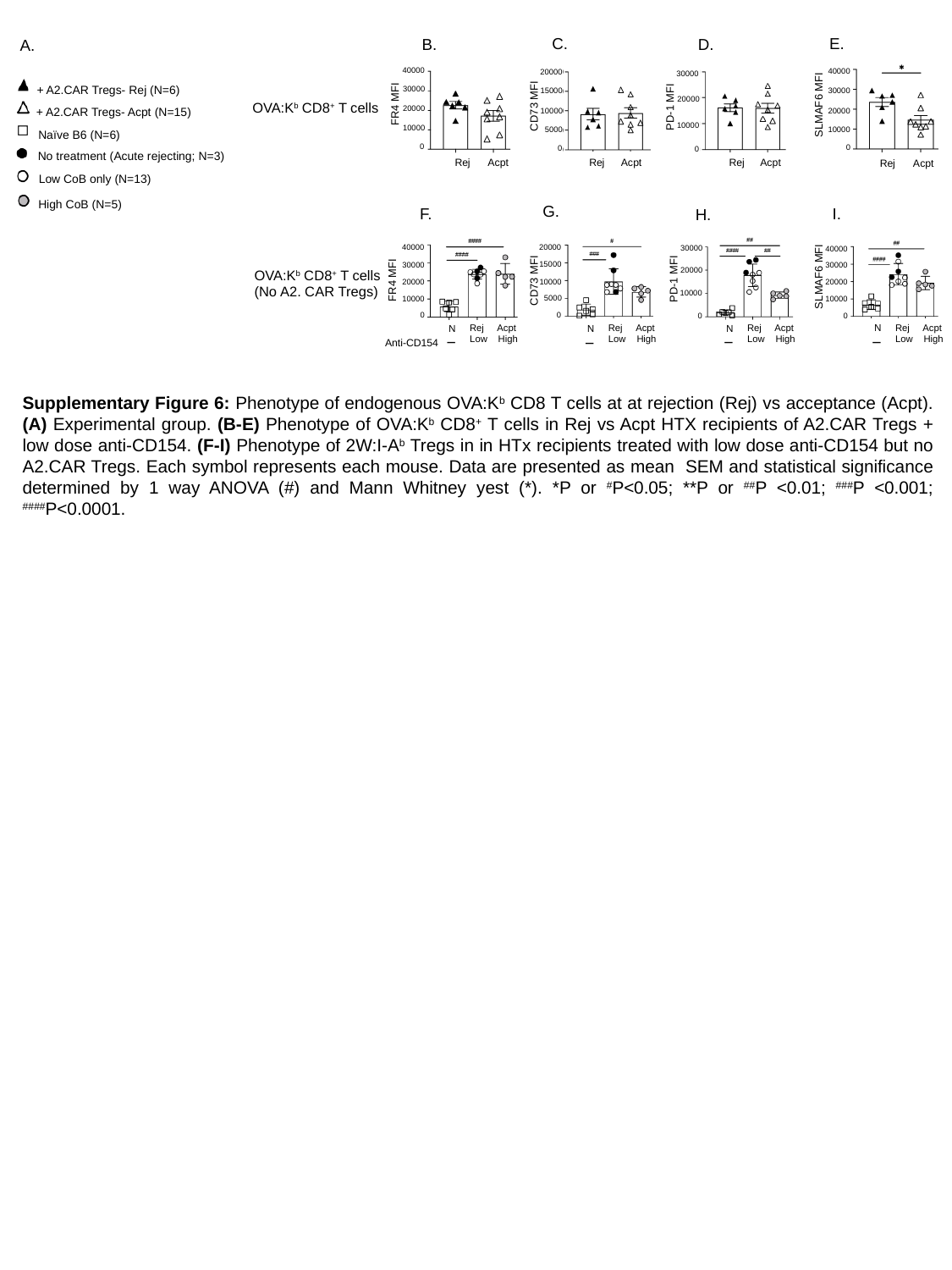

E.
C.
B.
D.
A.
40000
40000
20000
30000
+ A2.CAR Tregs- Rej (N=6)
30000
15000
30000
20000
FR4 MFI
SLMAF6 MFI
OVA:K^b^ CD8^+^ T cells
CD73 MFI
PD-1 MFI
20000
+ A2.CAR Tregs- Acpt (N=15)
20000
10000
10000
10000
10000
5000
Naïve B6 (N=6)
0
0
0
0
No treatment (Acute rejecting; N=3)
Rej
Acpt
Acpt
Rej
Rej
Acpt
Rej
Acpt
Low CoB only (N=13)
High CoB (N=5)
G.
F.
I.
H.
40000
20000
30000
40000
15000
30000
30000
20000
OVA:K^b^ CD8^+^ T cells
(No A2. CAR Tregs)
SLMAF6 MFI
PD-1 MFI
FR4 MFI
CD73 MFI
20000
20000
10000
10000
5000
10000
10000
0
0
0
0
Rej Acpt Low High
Rej Acpt Low High
N
Rej Acpt Low High
Rej Acpt Low High
N
N
_
N
_
_
_
Anti-CD154
